# Magnetic Fields Influence Visual Responses in Mice

**DOI:** 10.1101/2025.05.12.653455

**Authors:** Ma’ayan Semo, Steven Hughes, Nicola J. Smyllie, Andrew P. Patton, Carina A. Pothecary, S.K. Eric Tam, Julia Buckland, Laurence A. Brown, P. Kirk Reardon, David M. Bannerman, Mark W. Hankins, Michael H. Hastings, Stuart N. Peirson

## Abstract

Many animals use the Earth’s magnetic field for the purposes of orientation and navigation, although the sensory mechanisms remain unclear. It has been proposed that retinal responses to light may be modulated by magnetic fields. However, to date, there is no evidence for a retinal response to magnetic fields in mammals. Here we show that magnetic fields affect expression of the neuronal activity marker c-Fos in the mouse retina in a light dependent manner. These retinal responses to magnetic fields are abolished in mice lacking the candidate magnetoreceptor cryptochrome. To characterise the signalling pathways involved, we then used RNAseq and cell-type mapping. We also show that magnetic fields increase exploratory behaviour in a visually dependent task and lengthen the period of the retinal circadian clock. Together, our data provide the first evidence for a mammalian retinal response to magnetic fields at a cellular, molecular and functional level, which may influence vision.

## INTRODUCTION

Many species of animal are capable of detecting geomagnetic fields, which are used for orientation and navigation, particularly for the purpose of migration (*1*). However, the biological mechanisms underlying magnetoreception remain unclear (*2, 3*). It has been proposed that light-dependent radical pair intermediates may be capable of responding to magnetic fields, enabling magnetic information to be transduced to the sensory nervous system (*4*). The strongest candidate for this radical pair magnetoreceptor is the flavoprotein cryptochrome (CRY), which possesses the necessary chemical properties to be capable of responding to magnetic fields (*5*). CRYs are related to photolyases which repair DNA damaged by UV light, and also play a key role in the negative limb of the intracellular circadian clock (*6, 7*).

Animal CRYs occurs in two forms, type 1 CRYs, which acts as blue-light sensitive circadian photoreceptors, and type 2 CRYs, which function as a negative regulator of the core circadian clock mechanism and are not thought to be light sensitive (*8*). In *Drosophila*, a type 1 CRY plays a key role in light input to the circadian clock (*9, 10*). Studies in *Drosophila* have also shown that CRY plays a critical role in light-dependent detection of magnetic fields. This includes binary-choice behavioural assays (*11, 12*), negative geotaxis (*13*) and period-lengthening in constant light (*13, 14*). Genetic studies in flies have shown that the magnetic field responses are dependent on photosensitive type 1 CRYs (*11, 14*). However, both type 1 and type 2 insect CRYs can restore light-dependent magnetic field responses (*12*). Most surprisingly, human CRY2 can also restore magnetic field responses in CRY-null flies (*15*).

Amongst vertebrates, birds are known to be capable of responding to the direction of a magnetic field, which they use for navigation and seasonal migration (*5*). Studies in birds have shown that cellular markers of neuronal activity - such as c-Fos - are elevated in CRY-expressing cells during magnetic orientation. These cells primarily appear to be retinal ganglion cells, including displaced ganglion cells in the inner nuclear layer (*16*). However, it is unclear if this response is CRY-dependent. Recent studies have suggested that CRY4 may be the most likely candidate for the light-dependent magnetic compass in birds (*17–20*). By contrast, evidence for mammalian responses to magnetic fields is more limited. Mice have been shown to display a magnetic field response in spatial memory tasks (*21, 22*), nest building (*23*) and learnt orienting responses in a 4-arm water maze (*24*). Moreover, human visual sensitivity has also been suggested to be influenced by geomagnetic fields (*25, 26*). It has been proposed that rather than an independent magnetic sense, magnetic fields may influence visual spatial perception, providing additional information for navigation (*27, 28*). Together, these studies provide evidence that magnetoreception may occur in mammals but provide no evidence for the mechanistic basis of these responses. Moreover, the putative avian magnetoreceptor *Cry4* is only found in non-mammalian vertebrates (*29*).

In mammals, light input to the circadian system is dependent on retinal photoreceptors, including the classical rod and cone photoreceptors, as well as a subset of photosensitive retinal ganglion cells (pRGCs) expressing the photopigment melanopsin (*30–32*). Whilst CRY was originally proposed as a mammalian circadian photopigment, there is no evidence for a role of CRY in light input to the circadian system (*33–35*). However, the mammalian retina does express CRYs, which play a key role in regulating retinal circadian physiology (*36*). In mammals, two copies of the negative regulator type 2 CRY (but no type 1) are found, termed CRY1 and CRY2 (*37*). CRY1 is expressed in cone photoreceptors as well as amacrine and ganglion cells of the inner retina, whereas CRY2 is expressed throughout the photoreceptor layer. CRY1 appears to play a key role in molecular and physiological rhythms, including clock gene rhythms as well as rhythms in the photopic electroretinogram (ERG), contrast sensitivity and pupil constriction. By contrast, CRY2 appears to only play a role in rhythms in the cone ERG (*38*).

To determine if magnetic fields influence neuronal activity in the mammalian retina, here we studied retinal c-Fos expression in response to light and magnetic fields in mice. We show a clear magnetic field effect in c-Fos expression in the inner retina, which is elevated in darkness but attenuated in the light. We go on to show that this retinal response to magnetic fields is abolished in mice lacking CRY1 and CRY2. Using RNA sequencing we show that magnetic fields influence a significant number of retinal transcripts, with an enrichment of genes expressed in cones and amacrine cells. Finally, we show that magnetic fields increase exploratory behaviour during a visually dependent behavioural task and can lengthen the period of the retinal circadian clock – an effect that is inhibited by exposure to blue light. Together, our data provide the first unequivocal demonstration of a retinal response to magnetic fields in mammals, which is dependent on the putative magnetoreceptor, cryptochrome.

## RESULTS

### Retinal c-Fos induction in response to light

In the dark, c-Fos was primarily expressed in the photoreceptor layer. In response to light for one hour at ZT14, expression of c-Fos in the photoreceptor layer was reduced (**Fig S1A**). By contrast, in the inner retina exposure to light upregulated c-Fos expression (**Fig 1A**), as has been described previously (*39–45*). c-Fos induction occurred in retinal amacrine and ganglion cells (**Fig S1B**), including retinal ganglion cells expressing BRN3A and OPN4 (**Fig S1C-D**) and GABAergic amacrine cells labelled with GAD67 (**Fig S1E**). Expression was limited to amacrine and ganglion cells, with no colocalization in bipolar cells (labelled with CHX10) (**Fig S1F**). Due to the gradient in short-wavelength sensitive (S) cone opsin and middle-wave sensitive (M) cone opsin in the mouse retina, the level of c-Fos may depend upon the area of the retina sampled (*46*). To account for this gradient of cone opsin expression, we used the gradient of UVS opsin to orient each retina and counted cells in multiple regions of both upper (dorsal) and lower (ventral) regions of wholemount retina (**Fig S2A**). As we used a cool-white LED light source with little UV output, fewer c-Fos positive cells in the lower retina is expected given the predominance of UV sensitive cones in this region (*46*). As expected, due to this gradient of cone opsin expression, c-Fos responses were attenuated in the lower retina (**Fig S2B-C**). However, as similar effects of both light and EMF were observed in both upper and lower regions of the retina, total retinal c-Fos was used for all subsequent analysis.

**Fig. 1.**
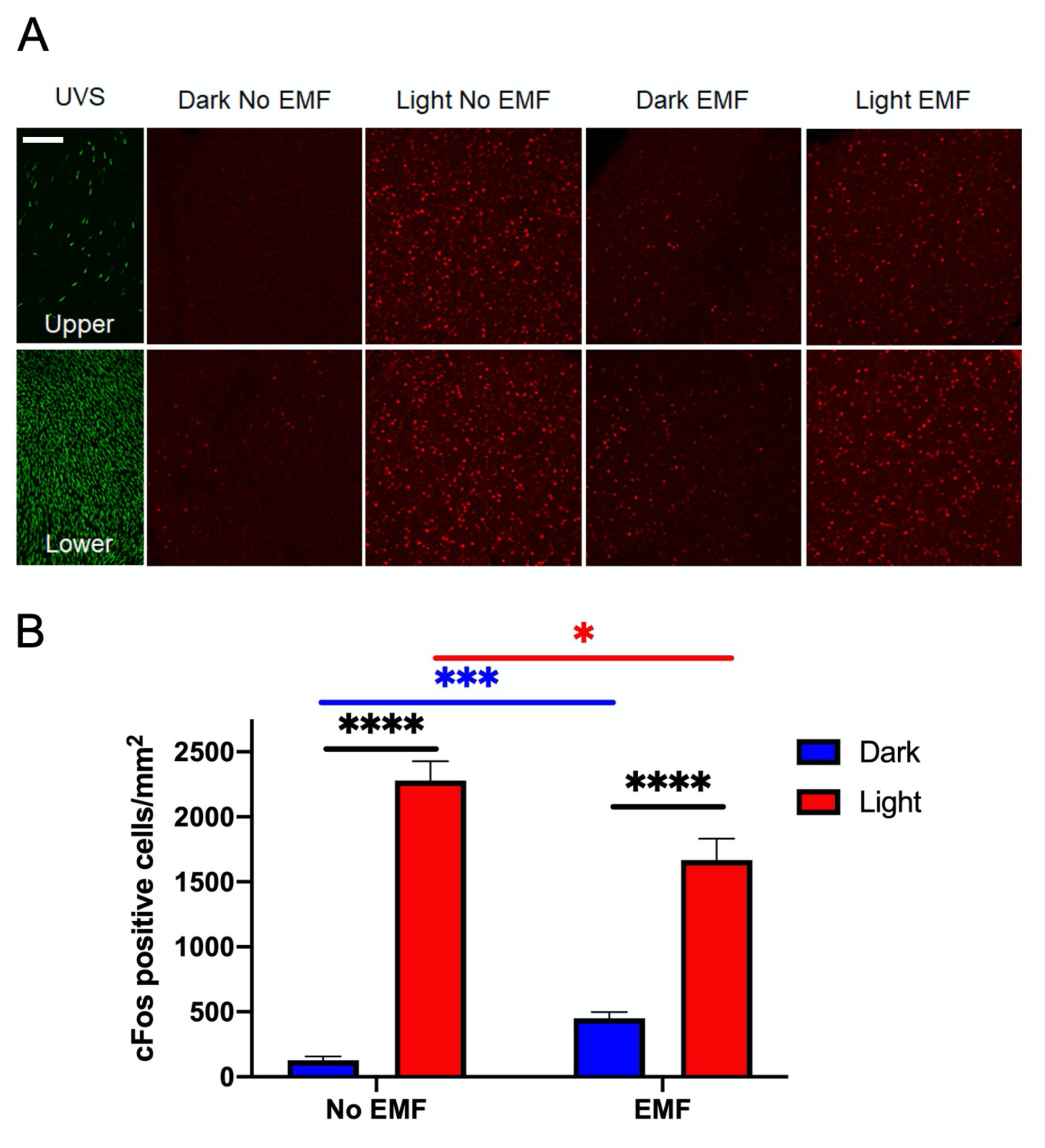
Retinal c-Fos induction in response to light and modulation by magnetic fields. Images of the distribution of c-Fos (red) and UVS opsin (green) in whole retina flat mounts from C57BL/6 wildtype mice. **(A)** Higher levels of c-Fos are induced by light (a cool-white LED light source with little UV output) in the upper retina (top row) than in the lower retina (bottom row). Light/Dark, EMF/No EMF condition are indicated above each column of images. Scale bar 100µm. **(B)** Quantification of c-Fos shows that EMF modulates the magnitude of c-Fos induction. Values plotted are means ±SEM with * indicating significance level (*Post Hoc* Bonferroni’s multiple comparison test): \**p*<0.05, \*\*\**p*<0.0001, \*\*\*\**p*<0.00001.

### Magnetic fields affect neuronal responses in the mouse retina

In the retina of mice exposed to a 100μT magnetic field for one hour immediately before tissue collection, increased c-Fos was observed in the dark. By contrast, in the light, reduced c-Fos induction occurred (**Fig 1A**). Data were analysed using 2-way ANOVA, with a main effect of light (F_1,38_=324.7, p<0.0001), no main effect of EMF (F_1,38_=1.036., p=0.3151) but a light x EMF interaction (F_1,38_=31.08, p<0.0001). Under control conditions, light resulted in a significant increase in cells expressing c-Fos as expected (p<0.0001). Similar light- induced c-Fos was seen when mice were exposed to a 100μT magnetic field (p<0.0001). However, in comparison with no EMF conditions, significantly elevated c-Fos levels were observed following EMF exposure in the dark (p<0.001), whereas decreased c-Fos occurred in response to light (p<0.05). As such, magnetic fields appear to increase neuronal activity in the dark-adapted retina, whilst attenuating responses to light.

### Retinal responses to magnetic fields are CRY-dependent

To investigate the role of cryptochromes in the retinal c-Fos response to magnetic fields, we then studied retinal c- Fos responses in mice lacking both Cry1 and Cry2 (*Cry1/2^-/-^*). A similar pattern of c-Fos induction in response to light was immediately apparent from retinal wholemount images (**Fig 2A, Fig S2C**). In mice lacking cryptochromes, there was a main effect of light (F_1,20_=276.5, p<0.0001), no effect of EMF (F_1,20_=0.1783, p=0.6773) and no light x EMF interaction (F_1,20_=0.5249, p=0.4771). Whilst the response to light was clearly preserved in mice lacking CRYs, no modulation of this response by EMF was apparent (**Fig 2B**).

**Fig. 2.**
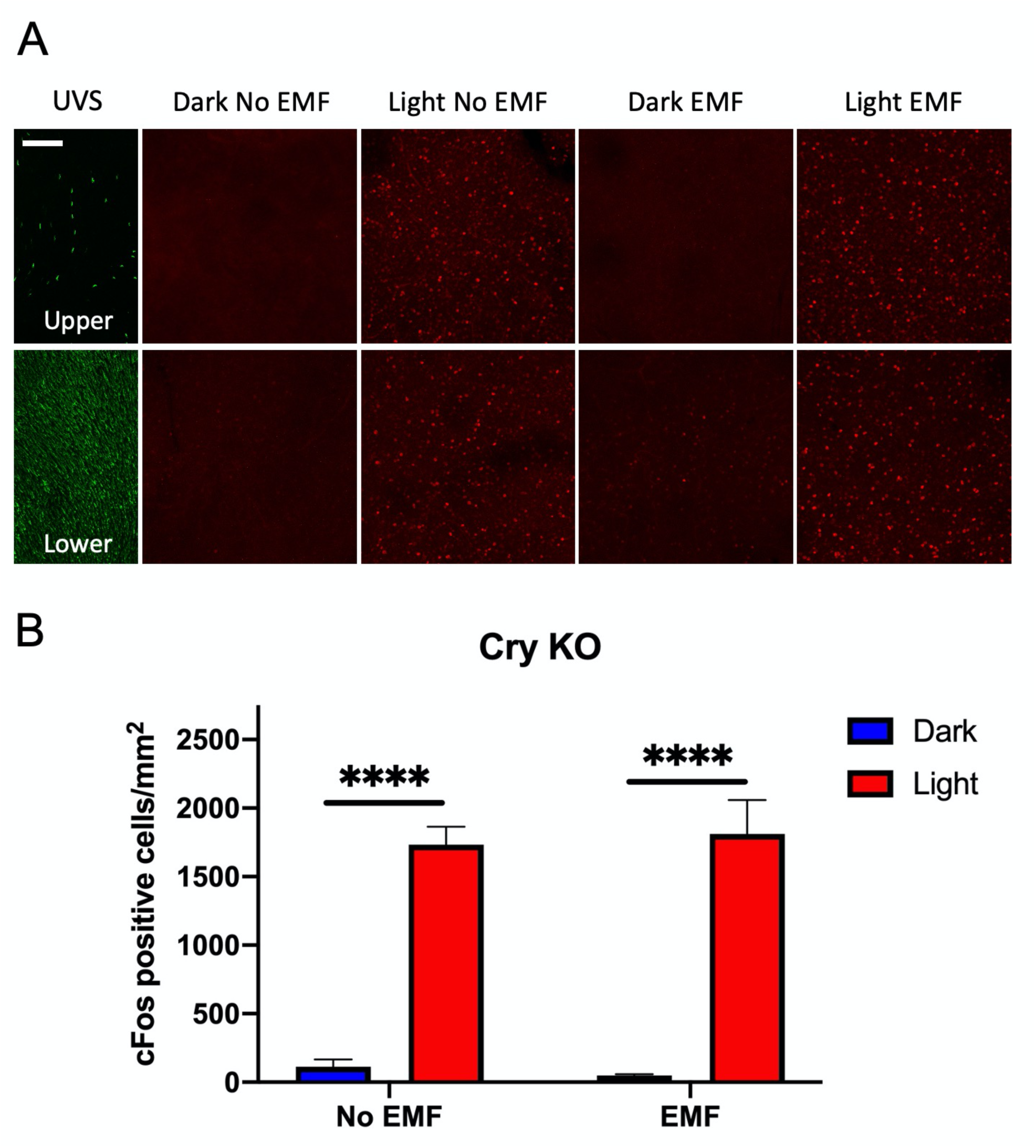
Modulation of retinal c-Fos induction in response to magnetic fields is abolished in CRY-deficient mice. Images of the distribution of c-Fos (red) and UVS opsin (green) in whole retina flat mounts from CRY-deficient mice (*Cry1^-/-^/Cry2^-/-^*) **(A).** The UV cone distribution is maintained in CRY-deficient mice. There are higher levels of c-Fos induced by light (a cool-white LED light source with little UV output) in the upper retina (top row) than in the lower retina (bottom row). Scale bar 100µm. **(B)** The magnitude of c-Fos induction is not modulated by EMF exposure in CRY-deficient mice. Values plotted are means ±SEM with * indicating significance level (*Post Hoc* Bonferroni’s multiple comparison test): \*\*\*\**p*<0.00001.

Retinal clock gene expression has recently been shown to influence the development of the UV sensitive opsin gradient in the retina (*47*). Whilst there was no loss of UV sensitive opsin gradient in CRY-deficient mice (**Fig 2A, Fig S2C**), closer inspection of retinal wholemounts showed a slightly reduced level of UV opsin expression in mice lacking CRY (**Figs S3A-B**). At higher magnification, some minor deficits in the organisation of the cone outer segments were apparent (**Figs S3C-D**). Given the comparable numbers of c-Fos positive cells in response to light in CRY-deficient mice, the loss of responses to magnetic fields do not appear to originate from overt developmental or structural changes in the retina. This is supported by data on visual responses in CRY-deficient mice (*38*).

### Retinal transcriptional responses to magnetic fields are distinct from light responses

Changes in retinal c-Fos immunoreactivity in amacrine and ganglion cells reflect a downstream consequence of light signalling, which originates with absorption of light by the rod and cone photoreceptors, as well as melanopsin-expressing pRGCs. As such, altered c-Fos responses to magnetic fields do not identify the origin of the magnetic field response in the retina or other cellular and molecular mechanisms involved in this response. To characterise better the retinal signalling mechanisms involved in responses to magnetic fields, we collected dark and light exposed retina under control and EMF conditions and used RNAseq to characterise the transcriptional responses to light. Differentially expressed genes were identified using 2-way ANOVAs using EdgeR (*48*) to reveal main effects of light, EMF and a light x EMF interaction for each gene. 798 genes showed a statistically significant (FDR<0.05) response to light, 287 genes showed a response to EMF and 103 genes showed a light x EMF interaction (**Fig 3A**). The transcriptional response of the retina to EMF was therefore distinct from that to light. Out of 798 genes responding to light, only 10 also showed a response to magnetic fields (*Cav2*, *Copg1*, *Cul3*, *Ehd4*, *Prrt2*, *Rabggtb*, *Rd3*, *Tbc1d30*, *Vps13c*, *A930004D18Rik*).

**Fig. 3.**
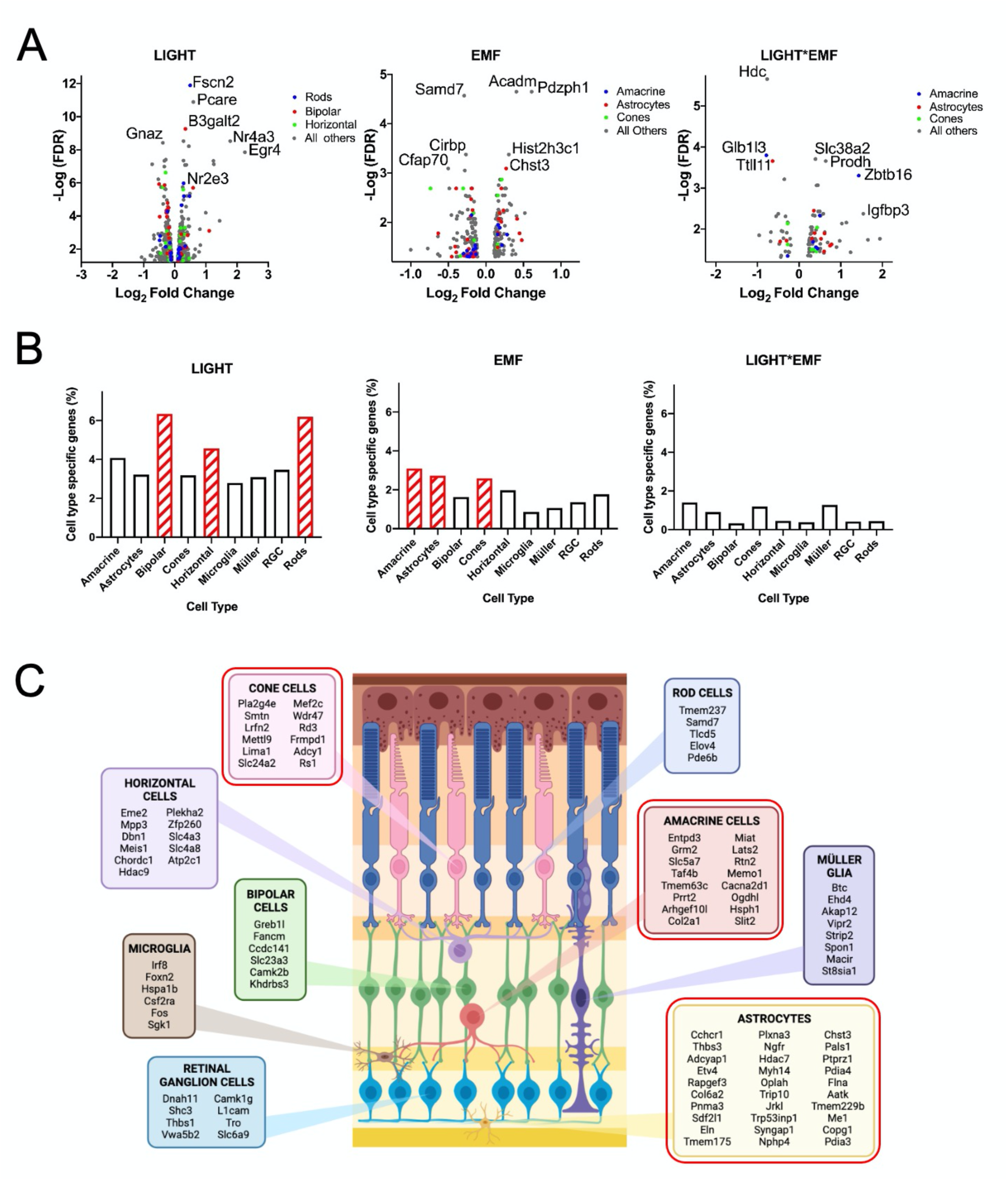
Characterisation of retinal gene expression in response to light and magnetic fields, and mapping of cellular responses. **(A)** Volcano plots showing the genes that are significantly changed in expression in response to main factors: light, EMF and Light x EMF identified using 2-way ANOVAs**. (B)** Proportion of retinal cell type specific genes in response to factors light, EMF and Light x EMF, the cell types that are significantly enriched by Fisher’s exact test (P<0.05) are indicated with red diagonal shading. **(C)** A schematic diagram of cell types in a section of retina, all the differentially expressed genes that map to a specific retinal cell type in response to EMF are indicated.

### Cellular origins of light and magnetic field responses

To understand the cellular origin of the retinal response to magnetic fields, we used published single-cell RNAseq data to map differentially expressed genes to specific cell types (**Fig 3B**). Whilst not all genes mapped to a specific retinal cell type, for light 236 out of 798 genes were mapped (30%); for EMF 102 out of 287 genes were mapped (36%); and for light x EMF conditions, 47 out of 103 genes were mapped (46%). In the light condition, there was a significant enrichment of genes (Fisher’s exact test) associated with rods (p=0.0132), horizontal cells (p=0.0186) and bipolar cells (p=0.0034). By contrast, in response to EMF, there was a significant enrichment of genes associated with cones (p=0.0360), amacrine cells (p=0.0160) and astrocytes (p=0.0010). No significant cell-type enrichment was apparent for light x EMF genes, likely due to the smaller number of genes involved. The differentially expressed genes mapped to each retinal cell type are shown for light (**Fig S4A**), EMF (**Fig 3C**) and light x EMF (**Fig S4B**).

### Amacrine cells expressing GRM2 show a distinct response to magnetic fields

The significant enrichment of genes associated with amacrine cells was of particular interest given the previously observed c-Fos response to light in this cell type. Amongst the genes responding to magnetic field exposure was *Grm2*, encoding type 2 metabotropic glutamate receptors. In the retina, GRM2 is expressed in starburst amacrine cells. To investigate if these cells showed a specific response to magnetic fields, we studied c-Fos in GRM2 expressing amacrine cells (**Fig 4A**). In the dark there is very little c-Fos expression within the inner retina; in EMF Off/Dark, the majority of the c-Fos cells that are present are also GRM2 positive (15 out of 19 cells, ∼79%). In the light there are many more c-Fos positive nuclei in the inner retina, however the proportion of c-Fos cells that are GRM2 positive is much reduced (in EMF off/Light, 17 out of 108 cells ∼16% are co-localised). When EMF is on in the dark there are more c-Fos cells present and the proportion colocalised with Grm2 is more similar to that of the light condition without EMF (in EMF On/Dark, 8 out of 40 cells ∼20% are colocalised). This is a significantly different proportion to that observed in the control condition EMF Off/Dark (Chi squared P<0.0001). EMF does not have a significant effect on proportions of c-Fos cells that are Grm2 positive when the light is on (in EMF On/Light, 45 out of 338 cells, ∼13% are colocalised) (**Fig 4B)**.

**Fig. 4.**
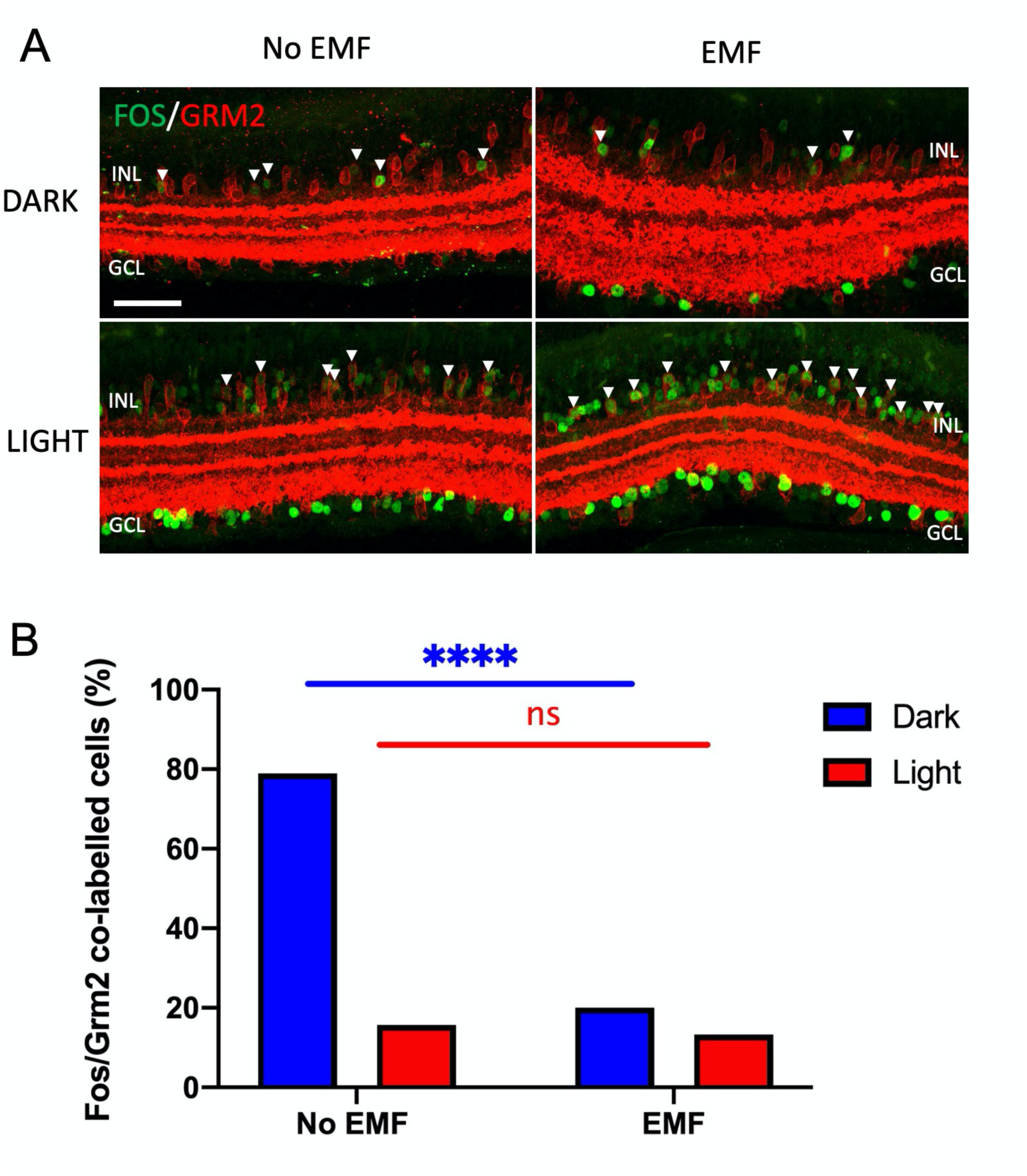
Modulation of starburst amacrine cell activity in response to magnetic fields. **(A)** Representative images of sections of retina stained for GRM2 (red) and c-Fos (green). The right-hand images are from animals exposed to EMF and those on the left not exposed, the tissues in the bottom row were exposed to light whilst the top row were in darkness. c-Fos cells that are positive for Grm2 are indicated with white arrowheads. Scale bar 50µm. **(B)** Bar chart showing the proportion of c-Fos cells that are co-labelled with GRM2. In the dark EMF condition, the proportion is significantly reduced compared to no EMF in the dark. In the light EMF exposure does not significantly change the proportion. Chi squared analysis, * indicating significance level: ***p<0.0001; ns, not significant.

### Characterisation of visually dependent behavioural response to magnetic fields

Starburst amacrine cells are known to play a role in direction selectivity, with targeted ablation of these cells impairing optokinetic nystagmus (*49*). To investigate if the optokinetic response was affected in response to magnetic fields, we studied both visual acuity (**Fig 5A**) and contrast sensitivity (**Fig 5B**) using the optomotor response in the presence and absence of magnetic fields. Whilst a subtle reduction in visual acuity was apparent, no significant differences were detected between control and EMF-exposed animals. Data were analysed using repeated measures two-way ANOVA. For visual acuity the only significant factor was cycles per degree (F_3,45_=95.32, p<0.0001), with most tracking occurring at 0.2 cycles per degree, whilst EMF (F_1,15_=0.8188, p=0.3798), and cycles/degree x EMF (F_3,45_=0.2179, p=0.8835) were not significant. For contrast the only significant factor was contrast (F_5,70_=89.44, p<0.0001) with an increase in tracking with increasing contrast, whilst EMF (F_1,14_=0.2465, p=0.6273), and cycles/degree x EMF (F_5,70_=0.3326, p=0.8916) were not significant.

**Fig. 5.**
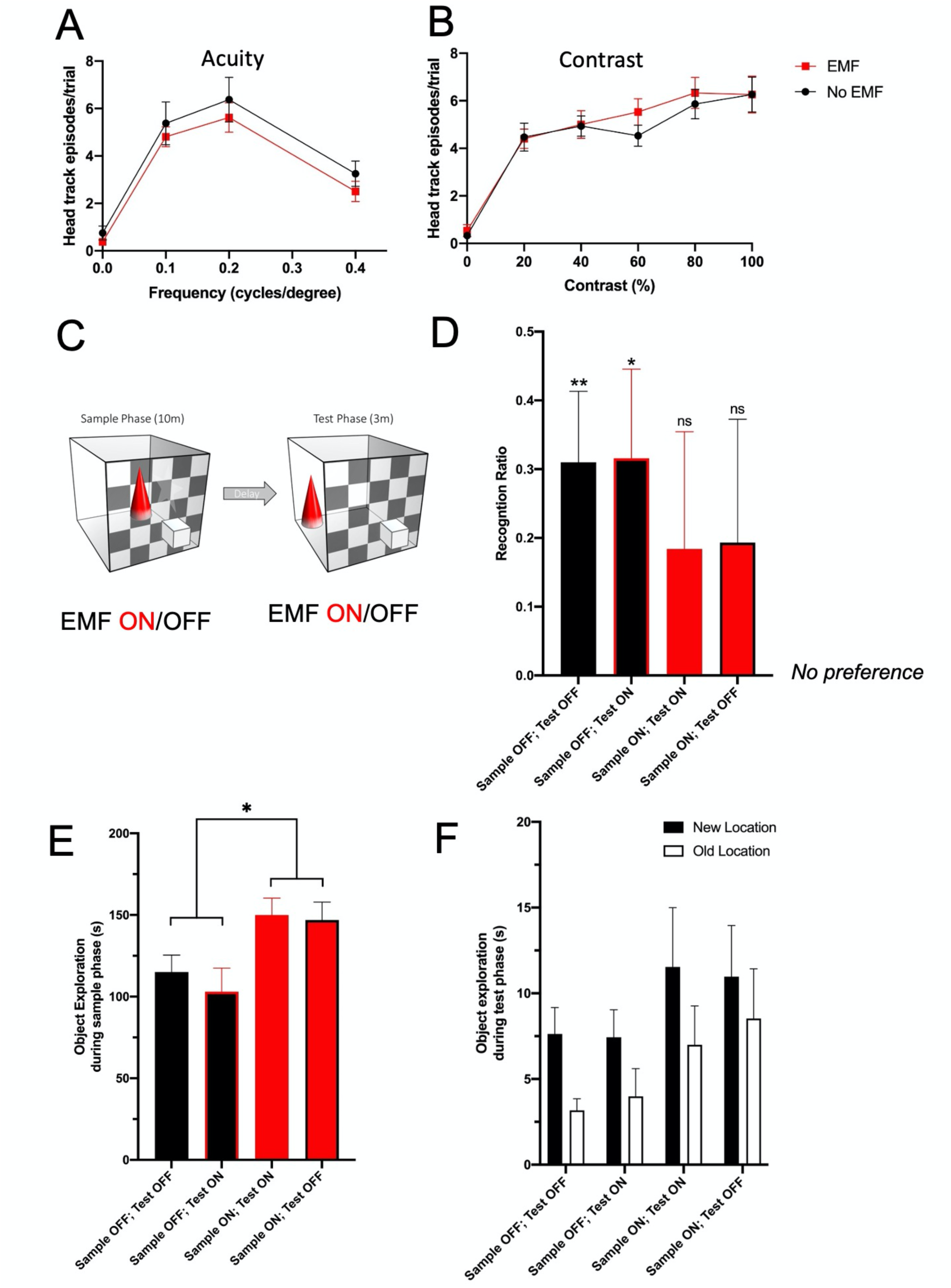
Exposure to magnetic fields does not significantly impair visually guided behaviours. Head tracking episodes in the presence (red) and absence (black) of EMF measured at different grating frequencies **(A)** and contrasts **(B)**. The number of episodes is not significantly altered under EMF exposure. **(C)** Schematic for the spatial version of the novel object recognition task. In the test phase animals were placed in an arena with visually distinct patterns on the walls and two different objects for 10 minutes. The animal was removed for 5 minutes during which time one of the two objects was moved, the animal was returned for the test phase of 3 minutes, in the presence or absence of EMF. **(D)** The object recognition ratios are above chance in both the absence and presence of EMF during the test phase when EMF is off in the sample phase (black bars). However, the object recognition ratios are not significantly above chance when EMF is on in the sample phase (red bars), regardless of whether EMF is on or off in the test phase. Significance indicated by * (Student’s t-test). **(E)** Object exploration in the sample phase of the test is significantly higher in the presence of EMF (red fill) than when it is off (black fill). Significance indicated by * (Two-way ANOVA: significant effect of EMF in sample phase)**. (F)** Object exploration during the test phase. There is significantly more exploration of the object in the new location, and more exploration following EMF exposure in the sample phase (three-way ANOVA: significant effects of location (new/old) and EMF in the sample phase). Error bars denote standard error of the mean. * *P*<0.05.

We have previously shown that the object displacement task in mice is visually dependent (*50, 51*). Moreover, recognition memory is also sensitive to changes in context, where changes in environmental conditions between the sample and test phase result in impaired performance (*51*). To test if magnetic field exposure affected visually dependent behaviour, we exposed mice to magnetic fields in either the sample and/or test phase in a counterbalanced manner (**Fig 5C**). We predicted that a change in EMF would impair performance on this visually dependent task – but only if the animals were able to detect this. Under control conditions (no EMF either phase), mice performed above chance (p = 0.0089 (2-tailed one sample *t*-test)). When mice were exposed to EMF during the test phase, they continued to perform above chance (p = 0.0278 (2-tailed)). However, when exposed to EMF during the sample phase – or sample and test phases - performance was not significantly above chance. Whilst performance was lower, the main effect of sample phase EMF exposure on spatial recognition was not statistically significant (p = 0.068, **Fig 5D**). Performance may be influenced by the time an animal spends exploring objects in the sample and test phases or locomotor activity, rather than any impairment in recognition memory. Whilst distance travelled was unaffected by EMF, mice exposed to EMF during the sample phase spent more time exploring objects in both sample AND test phases (main effect of sample EMF on object exploration during sample phase (**Fig 5E** F_1, 15_=7.484 p=0.015) and test phase (**Fig 5F** F_1,15_= 5.480, p = 0.033). These data reveal a subtle effect on exploratory activity of 10-min EMF exposure. This is consistent with previous findings on the impact of magnetic field on exploratory activity in laboratory mice and other rodent species (*52*). Previous studies have shown that *Cry1^−/−^, Cry2^−/−^* mice show low exploratory drive and deficits in recognition memory in novel object recognition tests (*53*). Due to these confounds in baseline behaviour, these studies were not performed in mice lacking cryptochromes.

### Effects on retinal circadian rhythms

Given the central role of cryptochromes within the circadian clock, we next tested the effect of magnetic fields on the clock mechanism within the retina. Bioluminescence rhythms were recorded from retinal explants of PER2::Luc mice and exposed to magnetic fields of strengths between 0 and 1200μT using a Schüderer apparatus-based system (**Fig 6A**). In these retinal explants, we saw effects on circadian period indicating that this tissue is sensitive to EMF treatment (repeated measures two-way ANOVA EMF/no EMF effect: F_1,62_=4.668, p=0.0346) and this was dependent on field strength (repeated measures two-way ANOVA: field strength effect: F_6,62_=2.576, p=0.0271; interaction: F_6,62_=4.040, p=0.0018). Interestingly, this effect of field strength was limited to a period lengthening at 800μT (Post-hoc Sidak’s multiple comparisons test: p<0.0001), but field strengths either below or above this strength had no effect on circadian period (**Fig 6B**). The period increase caused by 800μT field strength exposure was ∼0.7 hr compared with paired no EMF controls (**Fig 6C)** (paired two-tailed t-test: t_17_=5.334, p<0.0001). As 800μT field strength exposure was the only effective field strength tested, we focused on this for further investigation of the interaction between EMF exposure and diurnal light cycles. Due to the fact that light would be expected to entrain the retinal clock to a period of 24 hours, if EMF altered the intrinsic free-running period of the retina the relative phase of the entrained retinal oscillation would be altered. We assessed, therefore, the phase relationship between the rhythms of paired retina (control and EMF-exposed) in response to daily light cycles. We found that this was modulated in a wavelength-dependent manner (**Fig 6D**). Retina exposed to different lighting schedules exhibited different phase relationships dependent on EMF exposure (repeated measures two-way ANOVA light effect: F_2,33_=1.251, p=0.2995; EMF effect: F_1,33_=8.164, p=0.0073; Interaction: F_2,33_=2.986, p=0.0643). We therefore investigated the phase relationships in response to EMF within the different lighting regimes. As expected, PER2::Luc rhythms of retinas exposed to 800μT magnetic fields in the absence of light were phase delayed compared to no EMF conditions, due to period lengthening (Sidak’s multiple comparisons: p=0.0202). Similarly, retinas subjected to red (625nm) wavelength light cycles and magnetic field exposure were significantly phase-delayed relative to no EMF controls, that were also exposed to the same red light cycles (Sidak’s multiple comparisons: p=0.0212). In blue (405nm) wavelength light cycles, however, PER2::Luc oscillations remained phase-locked to the lighting cycle and were not modified by magnetic field exposure (Sidak’s multiple comparisons: p=0.9719). Thus, both blue light and EMF can influence circadian timekeeping in the retina, with blue light representing a stronger entraining signal.

**Fig. 6.**
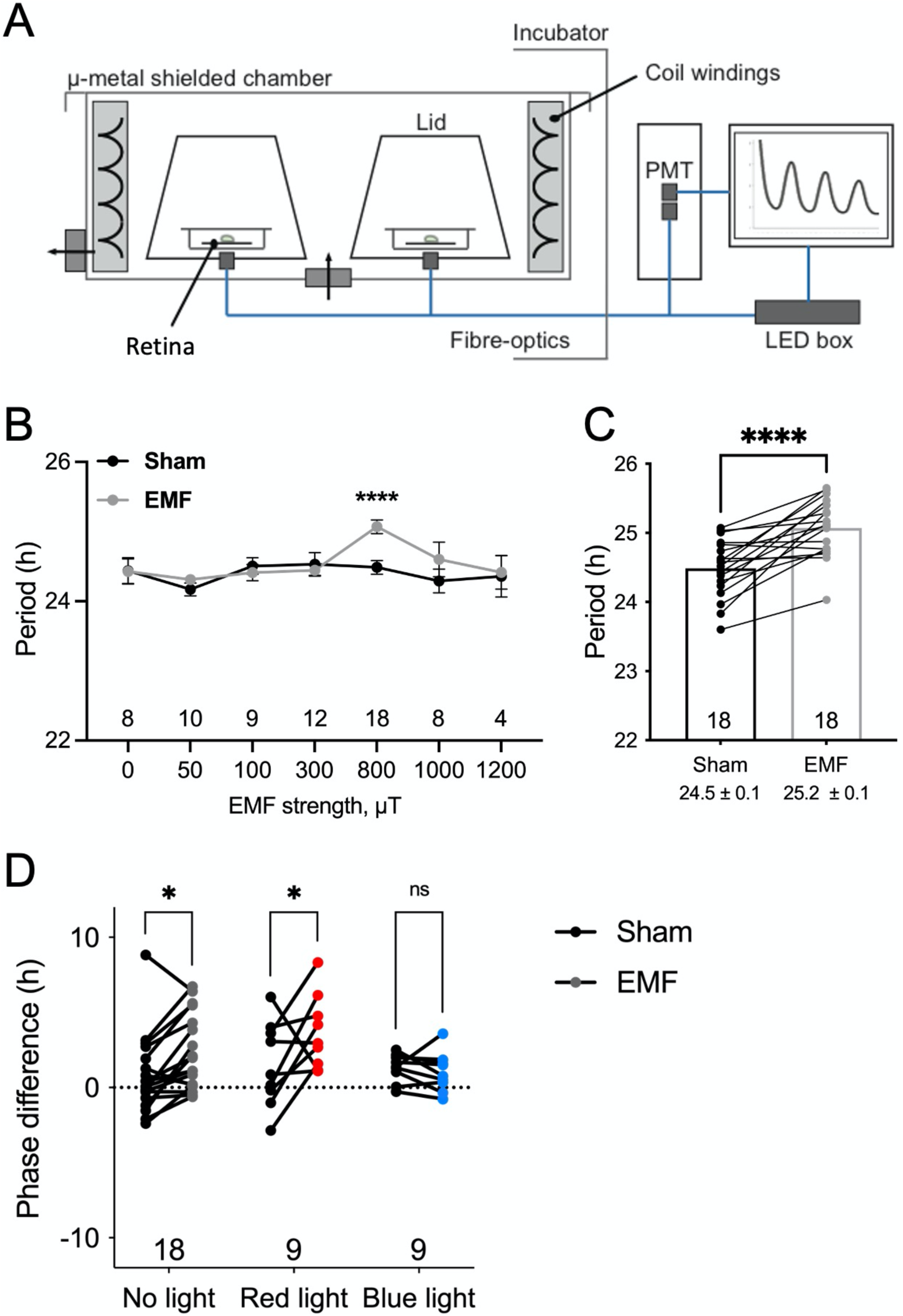
The effect of magnetic field exposure on retina circadian rhythms. **(A)** Schematic of the setup for *ex vivo* recording of retinal circadian rhythms of PER2::Luc bioluminescence. **(B)** Group data showing the effect of magnetic field exposure on retina explant circadian period at a range of field strengths from 0 – 1200μT. **(C)** A paired comparison of data in (B) showing significant circadian period lengthening of PER2::Luc rhythms in retinas exposed to 800 μT compared to no EMF controls. **(D)** Group data showing phase difference of PER2::Luc oscillation peaks at the start compared with peaks at the end of 7 days of magnetic field treatment. The peak phases were expressed relative to (predicted in the case of No light condition) lights off.

## DISCUSSION

Here we show for the first time a cellular, molecular and functional response of the mammalian retina to magnetic fields. Using expression of the immediate early gene c-Fos, we show that in darkness retinal c-Fos levels are elevated by magnetic fields, whereas c- Fos induction in response to light is attenuated by magnetic fields. Furthermore, the retinal responses to magnetic fields we describe in both dark and light conditions are abolished in mice lacking the putative magnetoreceptor cryptochrome, providing the first direct evidence for a role of cryptochrome in magnetoreception in mammals. Finally, exposure to EMF at 800 μT lengthened circadian period of the retina, and this effect was blocked by exposure to cycles of blue, but not red, light.

How can we reconcile the opposite responses to magnetic fields that we observe in light versus dark? One potential explanation may be the signal inversion that occurs during light signalling within the vertebrate retina. In the dark, photoreceptors are depolarised and release glutamate. In response to light, these cells hyperpolarise and reduce neurotransmitter release. By contrast, inner retinal neurones such as amacrine and retinal ganglion cells may either depolarise or hyperpolarise in response to light signals from the outer retina, corresponding to ON and OFF pathways (*54*). Whilst the neuronal response we describe occurs in the inner retina, this may not reflect the cellular origin of the response and may be a downstream event. As such, changes in c-Fos levels in the inner retina could plausibly originate from changes in signalling in the outer or inner retina. If magnetic fields influence cellular depolarisation – as has been shown in *Drosophila* (*55*) – a depolarising cellular response at the level of the photoreceptors would result in reduced light signalling and less c-Fos induction at the level of the inner retina. Conversely, direct effects at the level of depolarisation of inner retinal neurons would result in increased c- Fos. Finally, it is worth noting that depolarisation of inhibitory amacrine cells could also result in reduced c-Fos levels in other inner retinal cell types. The exact circuits involved in these responses clearly require further investigation.

Whilst the attenuated light response we describe may be consistent with CRY photochemistry, the increased c-Fos response in darkness is not. To function as a magnetoreceptor, CRY is thought to require light input to produce the free-radical triplet states which are necessary for magnetic field responses (*4, 5*). However, this mechanism is based on CRY undergoing a redox cycle involving photo-reduction and light- independent re-oxidation. Studies in which radical pairs mediate magnetic field responses in birds suggest that responses may occur in the dark and involve re-oxidation (*56*). More recently, the canonical CRY-dependent radical pair mechanism of animal magnetoreception has been challenged by findings in *Drosophila*, where expression of just the C-terminal 52 amino acids of dmCRY is sufficient to support magnetoreception at a cellular and whole organism level (*57*). These data are consistent with a radical pair mechanism, but as they are modulated by cytosolic availability of FAD, imply that redox reactions may play a key role. Radical pairs that are not photochemically generated can also contribute in this model, which may explain how magnetoreception may occur in darkness (*56*).

We should also emphasise that the dark conditions used in these studies cannot preclude very low light levels driving scotopic responses, including the dim red light used in animal husbandry and tissue collection. The retina is exquisitely sensitive to light, with rod photoreceptors capable of responding to single photons. This may be comparable with night-migrating birds, which require light for the CRY photochemistry necessary to detect magnetic fields but migrate at night when light levels are low. As rod photoreceptors may be active under such low-light conditions, it has been suggested that the magnetic compass of such species may be located in the cones to avoid competition with visual transduction (*5*). As CRY1 is expressed in both the cones and inner retina, whereas CRY2 is expressed in rod and cone photoreceptor outer segments (*38, 58*), the localisation of CRY is consistent with this hypothesis. Mice have rod dominated retinae with cones accounting for 2-3% of photoreceptors (*59*). Moreover, mice do not possess the double cones that occur in the avian retina, which have been proposed as a putative site of magnetoreception in birds (*5, 17*).

It has been proposed that a long-lived radical pair reaction product may be capable of influencing isomerisation of 11-*cis* retinal and thus altering classical phototransduction (*28*), although this model is not consistent with canonical cryptochrome photochemistry (*5*). The distinct transcriptional responses we describe here may reflect mechanisms by which magnetic fields influence specific retinal cellular circuits to modify the processing of visual information. Based upon our findings, we would tentatively suggest that magnetoreceptive radical pair reaction products may differentially influence cone and downstream amacrine cell signalling. Protein interaction data from birds suggest that CRY may be capable of interacting with elements of the phototransduction cascade (*19*) or via actions on glutamatergic signalling (*18*). If this is indeed correct, it would suggest that retinal magnetoreception and visual signalling are intertwined, which would make characterisation of these pathways more challenging than if they were strictly separate. However, from a biological perspective, this would seem more probable than the evolution of an independent magnetic field sense.

Here we show an effect on exploratory behaviour in response to magnetic field exposure, although it is unclear if this is specific to vision. Whilst starburst amacrine cells have been implicated in retinal directional selectivity and optokinetic eye movements (*49*), we found no change in optokinetic responses to frequency or contrast. The changes in exploratory behaviour observed may relate to changes in visual cues, particularly as the animal moves through its environment. However, is unclear whether the visual stimuli we investigated here are appropriate, or whether magnetic fields influence specific aspects of visual function. Detailed cellular physiology may be required to detect subtle changes in cellular signalling, as well as to define which specific retinal circuits and visual responses are affected. Finally, it should be noted that mammalian responses to magnetic fields may be poorly developed in comparison with species in which magnetoreception is highly developed, such as migratory birds. As such, whilst the mouse retina may be capable of responding to magnetic fields, such information may be less relevant in a nocturnal rodent which may be more dependent on sensory modalities other than vision.

In addition, circadian responses at the level of the retina were detected at higher field strengths (800μT). These findings show that magnetic fields are capable of influencing retinal circadian physiology, which are known to modulate a wide range of cellular and physiological processes to the varying demands of the 24h day (*36*). The effect of EMF, however, was weaker than that of daily cycles of blue light, a potent entraining signal of the retinal clock. Nevertheless, a role of magnetoreception in regulating the retinal molecular clock provides another mechanism by which cryptochrome may modulate retinal physiology. The lack of responses at lower field strengths, which were effective *in vivo*, may relate to the *ex vivo* nature of the retinal preparations used, where the physiological and anatomical contexts are inevitably incomplete. The responses seen, however, confirm an intrinsic sensitivity of the mammalian retina to EMF.

In summary, here we show for the first time that the mammalian retina shows a cellular response to magnetic fields that is dependent on the candidate magnetoreceptor cryptochrome. Transcriptional responses to magnetic fields appear to localise to cones and amacrine cells, in contrast to the rod, horizontal and bipolar cells that are enriched in light responses. This includes a subset of directionally sensitive amacrine cells expressing GRM2. We also show functional effects on exploratory behaviour and retinal physiology at the level of the molecular circadian clock. Characterising the effects of magnetic fields on mammalian retinal physiology provides an important step in understanding how anthropogenic electromagnetic fields may affect biology.

## ACKNOWLEDGEMENTS

*Cry1^-/-^/Cry2^-/-^* mice were a kind gift from Prof Gijsbertus T J van der Horst (Erasmus MC). This work was funded by the Electric and Magnetic Fields Biological Research Council. SNP is also funded by the Biotechnology and Biological Sciences Research Council (BB/S015817/1, BB/X002357/1). SNP and RGF are funded by Wellcome (106174/Z/14/Z) and SNP and MWH are funded by the Medical Research Council (MR/S026266/1). MHH, NJS and APP are funded by the Medical Research Council (MC_U105170643).

## AUTHOR CONTRIBUTIONS

SNP, conceived and designed the study. SNP, MWH, MHH, DMB, SH, MS, SKET, NS, AP, CAP, LAB designed the experiments. SH, MS, NS, APP, CAP, SKET performed the experiments. MS, SH, NS, AP, CAP, SKET, JB, PKR, LAB analysed the data. SNP, SH, MS, SKET, MHH, NS wrote original draft of the manuscript. DMB, MWH reviewed the manuscript.

## STAR METHODS

### KEY RESOURCES TABLE

| Reagent or Resource | Source | Identifier |
| --- | --- | --- |
| **Antibodies** |  |  |
| Goat poly. anti-Brn3a | Santa Cruz Biotechnology | Cat# sc-31984,<br>RRID:AB_2167511 |
| Sheep poly. anti-Chx10 | Abcam | Cat# ab16141,<br>RRID:AB_302278 |
| Guinea pig poly anti-c-Fos | Synaptic Systems. | Cat# 226 005,<br>RRID:AB_2800522 |
| Rabbit mono. anti-c-Fos | Cell signalling | Cat# 14609,<br>RRID:AB_2798537 |
| Mouse mono. anti-GAD67 | Millipore | Cat# MAB5406,<br>RRID:AB_2278725 |
| Rabbit poly. anti-mGluR2 | Abcam | Cat# ab150387,<br>RRID:AB_2715600 |
| Rabbit poly. anti-Opn4 | Advanced Targeting Systems | Cat# UF006,<br>RRID:AB_2314781 |
| Goat poly. anti-UVS | Santa Cruz Biotechnology | Cat# sc-14363,<br>RRID:AB_2158332 |
| **Experimental models: organisms/strains** |  |  |
| C56BL/6 mice | Envigo | RRID: MGI:5811150 |
| *Cry1<sup>-/-</sup>/Cry2<sup>-/-</sup>* mice | G T J van der Horst (Erasmus MC) | RRID: MGI:2181205,<br>MGI:2181206 |
| *Per2::Luc* mice | Jax Laboratories | RRID:IMSR_JAX:006852 |
| **Software and Algorithms** |  |  |
| ImageJ software | https://imagej.net/ | RRID:SCR_003070 |
| Any-maze | Stoelting, Wood Dale, IL | RRID:SCR_014289 |
| GraphPad Prism | GraphPad Software LLC | RRID:SCR_002798 |
| HISAT2 | toolshed.g2.bx.psu.edu/repos/iuc/hisat2/hisat2/2.2.1+galaxy1 | RRID:SCR_015530 |
| edgeR | toolshed.g2.bx.psu.edu/repos/iuc/edgeR/edgeR/3.36.0+galaxy0 | RRID:SCR_012802 |
| featureCounts | toolshed.g2.bx.psu.edu/repos/iuc/featurecounts/featurecounts/2.0.3+galaxy1 | RRID:SCR_012919 |
| Biorender | http://biorender.com | RRID:SCR_018361 |

### EXPERIMENTAL MODEL AND SUBJECT DETAILS

#### Animals

All animals were housed under controlled 12:12 light:dark cycles in a light controlled chamber for at least 10 days prior to all studies. C57BL/6 mice (Envigo, UK) were used for all studies, with the exception of those on CRY-deficient mice, which used *Cry1^-/-^/Cry2^-/-^*mice (*7*). PER2::Luc (*60*) mice were used for retina bioluminescence recordings.

#### Ethical statement

All procedures were conducted in accordance with the UK Animals (Scientific Procedures) Act 1986, under PPLs PE4ED9D2C/PP0911346 and PILs IFA195639, IB2F9F14B, I869292DB, I18656989 and I99083249.

### METHOD DETAILS

#### Magnetic field exposure

For all exposure studies, animals were housed in a custom-made black circular Perspex exposure chamber. This chamber was divided into two separate housing zones, enabling two animals to be studied at a time (**Fig S5**). Each zone had a separate food hopper and water was provided in a ceramic bowl (both available *ad libitum*). Each zone was ventilated and lit by an independent light source with an irradiance of 100 lux (cool white LED, Quadica Development, Lethbridge, Canada). The housing chamber contained no metal components and fitted into a custom-made bifilar wound Helmholtz coil (diameter 50cm). When active, this coil generated a field strength of 100μT, or sham “no EMF” condition in which electricity was still passed through the coil but no magnetic field was generated. All studies were performed in the presence of the Earth’s magnetic field (30-50μT).

Male WT C57 BL/6 mice (10-12 weeks of age) were maintained group housed in light-tight chambers under 12:12 lighting conditions (cool white LED (LuxeonStar, Quadica Developments Inc, Alberta, Canada), 100 lux, lights on 01:00 – 13:00) for at least 2 weeks prior to experiments. On the day of experiments, mice were placed into the exposure chamber at ZT 10, at the end of their light phase and while lights were still on in their home chambers (the main room lights were also on). On each day, 6 mice were placed into the EMF exposure chamber at ZT10, with 3 mice placed into each side. Each side of the chamber contained sawdust covering the floor, a small amount of bedding material in one corner and a water bowl placed on the floor of the enclosure in a fixed location. Food was available *ad libitum* via the side gratings. The exposure chamber lights were controlled by electrical timers and maintained on the same light cycle as the colony chambers (lights on 01:00 – 13:00), as such the lights within the exposure chamber remain on for 2 hours after mice are placed into the arena (ZT10-ZT12). Light intensity within the chamber was 100 lux cool white LED (LuxeonStar, Quadica Developments Inc, Alberta, Canada) and in darkness, light levels were <0.1 lux. At ZT14, two hours after lights off, mice were subjected to different combinations of Light and EMF exposure (as described below) for 1 hour from ZT14 to ZT15. Light pulses were administered by manually pressing a button on the light timer at ZT14. EMF or no EMF exposure was administered by manually switching on the power supply unit controlling the EMF exposure system, with a separate toggle switch set to provide EMF or no EMF exposures. Following a 1-hour treatment, mice were removed from the EMF exposure chamber one at a time and immediately culled by cervical dislocation and tissue collected for ICC studies (one eye fixed in 4% PFA) or RNA analysis (retina dissected from one eye and flash frozen on dry ice). Protein and RNA samples were collected and processed using separate sets of dissection tools. For groups not involving a light pulse, the collection of tissue was performed under dim red light (>600nm, <10 lux, provided from overhead room lighting and a head-torch worn by the experimenter). For groups involving a light-pulse, tissue collection was performed with the main room white lights on. Neither light-pulses nor EMF exposures were switched off until tissue had been collected from all mice (i.e., all mice stayed under the same constant conditions with no ‘Lights OFF’ transition for example). The time for collecting all tissue from n=6 mice of each group was typically around 20 minutes.

#### Immunocytochemistry

Isolated whole eyes were punctured via the anterior aspect using a 25 gauge needle and then placed into microfuge tubes containing 1ml of freshly prepared 4% methanol-free PFA (Thermo Fisher) and stored overnight at 4°C, then washed briefly in PBS, before being placed into 30% sucrose (w/v) in PBS at 4°C for >48 hours. For ICC of whole retina flat-mounts, retinas were dissected from eye cups then placed into 24-well plates and immersed in sterile PBS containing 30% sucrose. Retinas were subjected to a freeze thaw cycle by placing the 24-well plate onto a surface of dry ice until frozen, and then thawed at room temperature. Whole retinas were then moved to fresh plates containing 1% Triton-X in PBS and further permeabilised for 30 minutes at room temperature. Retinas were blocked using 10% normal donkey serum (NDS) in PBS with 1% Triton-X for 60 minutes and then incubated with primary antibodies for 3 days at 4°C (rabbit anti-c-Fos, goat anti- Brn3a, goat anti-UVS opsin; see **Table 1**). Retinas were then washed with PBS containing 1% Triton-X for 30 mins x3 and then incubated with secondary antibodies overnight (*c-Fos*/UVS co-labelling donkey anti-rabbit Alexa568 and donkey anti-goat Alexa488 Invitrogen) diluted in PBS with 1% Triton-X and 2% NDS. Retinas were then washed with PBS containing 1% Triton-X for 30 mins x4 and then counterstained with 0.2µg/ml DAPI in PBS for 5 mins at room temperature, before radial cuts were made and the whole retina mounted onto a microscope slide using Prolong Gold anti-fade mounting media (Thermo Fisher).

**Table 1.** Antibodies used in this study.

| Antigen | Antiserum | Dilution | Source, Product Number |
| --- | --- | --- | --- |
| Brn3a | Goat poly. | 1:200 | Santa Cruz Biotech, sc-31984 |
| Chx10 | Sheep poly. | 1:500 | Abcam, ab16141 |
| c-Fos | Guinea pig poly. | 1:2000 | Synaptic Systems, 226005 |
| c-Fos | Rabbit mono. | 1:200 | Cell Signalling, 9F6 14609S |
| GAD67 | Mouse mono. | 1:250 | Merck, MAB5406 |
| mGluR2 | Rabbit poly. | 1:1000 | Abcam, ab150387 |
| Opn4 | Rabbit poly. | 1:2500 | Advanced Targeting Systems UF006 |
| UVS | Goat poly. | 1:500 | Santa Cruz Biotech, sc-14363 |

For staining of retinal sections, eyes were fixed in 4% PFA then cryo-protected as above then the cornea and lens were removed and the eyecups embedded in OCT and sectioned at 18µm onto poly-lysine coated slides. Sections were blocked using 5% NDS in PBS with 0.3% Triton-X for 60 minutes and then incubated with primary antibodies (guinea pig anti- c-Fos and rabbit anti-GRM2 (mGluR2), see **Table 1**) in 1%NDS, PBS with 0.3% Triton-X) overnight at room temperature. Sections were then washed with PBS containing 2% NDS 0.3% Triton-X for 30 mins x3 and then incubated with secondary antibodies for 2 hours at room temperature (donkey anti-Rabbit Cy3 and donkey anti-guinea pig Alexa 488 Jackson ImmunoResearch) then counterstained with 0.2µg/ml DAPI in PBS for 5 mins at room temperature and finally coverslipped using Vectashield.

#### c-Fos image collection and analysis

Confocal images were collected using a LSM 710 laser scanning confocal microscope and Zen 2009 image acquisition software (Zeiss). Individual channels were collected sequentially. Laser lines for excitation were 405 nm, 488 nm, and 561 nm. Emissions were collected between 440-480, 505-550, and 580-625 nm for blue, green and red fluorescence respectively. For each retina, 3 non-overlapping areas of the dorsal and ventral regions were imaged at low magnification (x10 objective, image size 0.531 x 0.531mm) to maximise the area covered and number of cells analysed. Orientation of the retina was determined by visualisation of the dorsoventral gradient in UVS cone opsin expression observed in the adult mouse retina. Image slices were collected at 2.5um intervals from the GCL to the INL (typically 9-10 stacks per image) and including all layers of the retina for which c-Fos expression was detected. To enable direct quantitative comparisons, all image acquisition settings (pin hole size, laser power, gains setting, pixel dwell times) were identical for all images collected from all retinas, with all images from a given experiment collected in a single confocal user session. All image stacks were converted into maximum image projections using Zen 2009 software (Zeiss) and saved as separate TIFF files.

Quantification of immunofluorescence levels in confocal images collected from whole retinal flat-mounts was performed using ImageJ software (*61*) (NIH; rsbweb.nih.gov/ij/). Image analysis was performed using default settings in ImageJ. In order to ensure all images were processed under identical conditions, all individual images for comparison were first compiled into a single image stack in Image J. Background subtraction was performed on all images within the stack using automated settings within ImageJ. The de- speckle function was then applied to all images in the stack, followed by automated threshold analysis using standard settings in image J (with minor manual adjustment of automated threshold values, applied equally to all images in the stack). For a first-pass analysis, the mean pixel intensity was calculated for each image and used as a quantitative measure of intensity of immunofluorescence signal (i.e., the equivalent of turning the image into a single pixel with a value ranging from 0-256), where increased levels of c-Fos equate to higher mean pixel intensity values. For more detailed analysis, the particle analysis feature of image J was used to identify individual c-Fos positive nuclei in each image. The analysis tool was set to allow detection of objects with holes and a minimum size of 20 pixels was set to prevent any remaining artefacts being counted. For all images in the stack, the number of c-Fos positive nuclei, the mean intensity of each nucleus, and the area of each nucleus detected above threshold was calculated automatically. Values were exported to as .csv for subsequent statistical analysis. For all images, global enhancement of brightness and contrast was performed using Zen Lite 2011 image analysis software (Zeiss).

Co-labelling for GRM2/c-Fos was assessed in sectioned retinas of animals that had been exposed to EMF, or No EMF, with or without the presence of light (EMF off/Dark n=3; EMF off/Light n=2; EMF on/Dark n=3; EMF on/Light n=4). Confocal images were taken in regions of central retina in sections containing the optic nerve head (317.44 x3167.44µm) using an Olympus Fluoview FV1000. c-Fos positive cells were assessed for co-labelling with GRM2 in confocal stacked images within Image J (Fiji). One image per animal was assessed, apart from in the group EMF off/Dark in which two were assessed due to the relatively few c-Fos positive cells to be found in this condition.

#### Transcriptomic tissue collection

After tissue collection, retina for RNAseq analysis were stored at -80°C. Frozen retinal tissue was homogenised in TRIzol Reagent (Thermo Fisher) and total RNA isolated using RNeasy spin columns (Qiagen) with on-column DNase treatment (Qiagen). All samples had RNA concentrations >50ng/ul and 260/280 ratios >1.8 measured by spectrophotometry (NanoDrop). Samples were then run on an Agilent Bioanalyser (Agilent) to assess RNA integrity, all giving RIN numbers >8. Poly(A) RNA sequencing was conducted on 500ng of total RNA diluted to 33ng/µl in ultrapure water and loaded on multi well plates, with samples from different groups evenly distributed across the plate. Strand-specific cDNA libraries were prepared using Illumina’s TruSeq Stranded Sample Prep Kit (125-bp single-read mode) following the manufacturer’s directions, then sequenced on a HiSeq 4000 (Illumina Inc., San Diego, CA, USA) at a read depth of 20 million reads. Sequencing was conducted by the Oxford Genomics Centre (www.ogc.ox.ac.uk).

#### Tests of visually guided behaviour

Optokinetic responses were performed using head tracking using black/white stimuli of different width to test visual acuity and different greyscale contrast to test contrast sensitivity. A physical visual tracking drum (*62*) (30cm internal diameter, 55cm high, with the mice on a platform 20cm high inside the drum), was placed inside the EMF coils. The drum was illuminated from above at 100 lux (LuxeonStar, Quadica Developments Inc, Alberta, Canada) (*51*), and the responses were recorded using a Logitech USB web camera and ANY-MAZE software (version 4.5; Stoelting, Wood Dale, IL). Each trial consisted of 2 minutes total rotation (30 sec. clockwise, 30 sec. counter-clockwise and then repeated once), and each mouse was tested with either EMF “on” or “off” condition. For acuity testing, square wave gratings of 0.1, 0.2, 0.4 cycles per degree (C/D) and a blank (white) were used. Contrast sensitivity was measured using a stripe pattern of 0.2C/D. In order to keep the overall brightness of the patterns consistent the grey levels of the darker stripes were reduced by the same amount that the lighter stripes were darkened (*63*). The percentage contrast quoted is the difference in the greyscale level between the dark and light stripe (contrasts tested: 80, 60, 40, 20 and 0% between light and dark stripes). The videos were coded and analysed by another scientist who was blinded as to the EMF condition. A bout was counted when the animal was attending to the grating and its head moved in the direction of the gratings by more than 15°.

To investigate if magnetic fields impact visually-dependent behaviour, we conducted a spatial recognition task. The animal is exposed to two objects during the sample phase (10 mins) and – after a 5 mins interval - one of these objects is moved to a new location relative to the other during the test phase (3 mins). The animal is expected to spend more time exploring the object in the new location. We have previously shown that this test is visually dependent. Spatial object recognition was performed as described previously (*50, 51*). In all the tests the person performing the behavioural test was blinded as to whether the coils were in the EMF “on” or “off” condition. Responses were recorded using a Logitech USB web camera and ANY-MAZE software (version 4.5; Stoelting, Wood Dale, IL).

#### Retina bioluminescence recordings

Whole-retina explants were prepared from adult (>30 days old) PER2::Luc mice from both sexes and mounted on to Millicell Culture Inserts (EMD Millipore). Retina explants were maintained in culture at 37°C, 5% CO2 for 24 hours after preparation andthen transferred to sealed 35mm dishes for luciferase bioluminescence recordings for ∼7 days, measured by photon multiplier tubes (Hamamatsu Photonics, Hamamatsu), as previously described (*64*). Photon multiplier tubes were housed in an incubated (37°C), light-tight Schüderer apparatus-based system, where magnetic field and light exposure were delivered as previously described (*13*). The effects of EMF on the retinal clock were determined in continuous darkness (i.e., free-running) or under entrained conditions. To achieve the latter, 405nm or 625nm wave-length light from LEDs (1μW/cm^2^) (Thorlabs) was delivered as part of a 12h light:12 dark schedule (chronic light exposure alone can disrupt the clock mechanism and so was not suitable for this study). PER2::Luc bioluminescence emissions were acquired every minute when lights were off. During the 12 hours of the “lights on” part of the cycle, light was delivered as “mini-cycles” of 20 minutes on followed by 10 minutes off to record readings of bioluminescence, thus 20, 1-minutereadings of bioluminescence were acquired every hour during “lights on”. The two retinas from each animal underwent EMF on or EMF off exposure and thus allowed for within-animal, paired comparisons.

### STATISTICAL ANALYSIS

#### Immunocytochemistry and tests of visually guided behaviour

Count data was transformed Y=SQRT(Y+0.5). All immunocytochemistry and tests of visually guided behaviour were analysed in Graphpad Prism 9, using the statistical test described in the results.

#### RNAseq data analysi*s*

Paired-end reads were assessed for quality using FastQC, showing all samples to be of high quality (phred scores >30), with an average read depth of 18.6 million mapped reads. Reads were aligned to the mouse reference genome (build GRCm38/mm10) using HISAT2 (*65*) and gene level expression was derived using Featurecounts (*66*). Analysis of differentially expressed genes was conducted using EdgeR (*48*) with an FDR cut-off of 0.05. A 2-way ANOVA was conducted across the sample groups to assess the main effect of light, the main effect of EMF and light x EMF interactions.

#### Mapping of differentially expressed genes to retinal cell types

Differentially expressed genes were then mapped to retinal cell types using published data from a single-cell transcriptome atlas of the retina (Lukowski et al., 2019). This study identified 18 clusters, comprised of 10 cell type categories (rods, cones, horizontal cells, bipolar cells, retinal ganglion cells, amacrine cells, Müller cells, astrocytes, microglia and others). Where multiple clusters corresponded to one cell type, data were averaged, weighted by cluster size. For each gene, the percentage of global retinal expression accounted for by each of the 10 cell type categories was then calculated. The maximum expression was then used to assign genes to specific cell types, with a minimum criterion that a cell type should account for at least 30% of all retinal expression. To ensure specificity to a single cell type and minimise false positives, an additional criterion was included, requiring the expression level of the second-place cell type to be less than 65% of that of the first-place cell type for each gene. At this level, 10,748 human genes could be assigned to a specific retinal cell type. The ‘others’ category was also excluded as it expressed a mixture of rod, cone, Müller cell and RPE markers, which was not functionally relevant. This approach was tested for 83 experimentally validated cell-specific markers identified from the literature, accurately assigning 88% of known markers to the correct cell type (Buckland et al., *manuscript in preparation*). Higher expression criteria increased specificity but resulted in a lower gene>cell mapping with increased false negatives, whereas lower expression criteria produced higher gene>cell mapping, but with an increase in false positives. When translated from human retina to genes expressed in the mouse retina this method was able to assign 6,007 mouse genes to a specific cell-type (manuscript in preparation). Over-representation of specific cell types, amongst those differentially expressed genes which mapped to single cell types according to this model, was then tested for using Fisher’s exact test. Networks of genes were subsequently mapped using Cytoscape 3.8 using the STRING app (*67*) and illustrated using Biorender (Biorender.com).

#### Retina bioluminescence circadian analysis

Retina PER2::Luc bioluminescence recordings were analysed within Prism (Graphpad). Raw data were first de-trended using a centred fifth-order polynomial before determining “peak-to-peak” circadian period of the oscillation. Mean ± SEM period was determined for each retinal explant from the peak-to-peak periods (average of 6 peak-to-peak intervals for each retina). The peak phase measures were calculated on a peak-to-peak basis, relative to “lights off” of the entrainment schedule (or predicted “lights off” for the no light condition). The difference between this relative phase measure at the start and the end of the recordings was defined as the phase difference after each treatment (light and/or EMF).

## SUPPLEMENTAL INFORMATION

Supplemental information includes 5 figures and supplementary data file “Raw Figure Data.xlsx”. Fastq files for RNAseq data are available from Gene Expression Omnibus https://www.ncbi.nlm.nih.gov/geo/

## SUPPLEMENTARY FIGURES

**Fig. S1.**
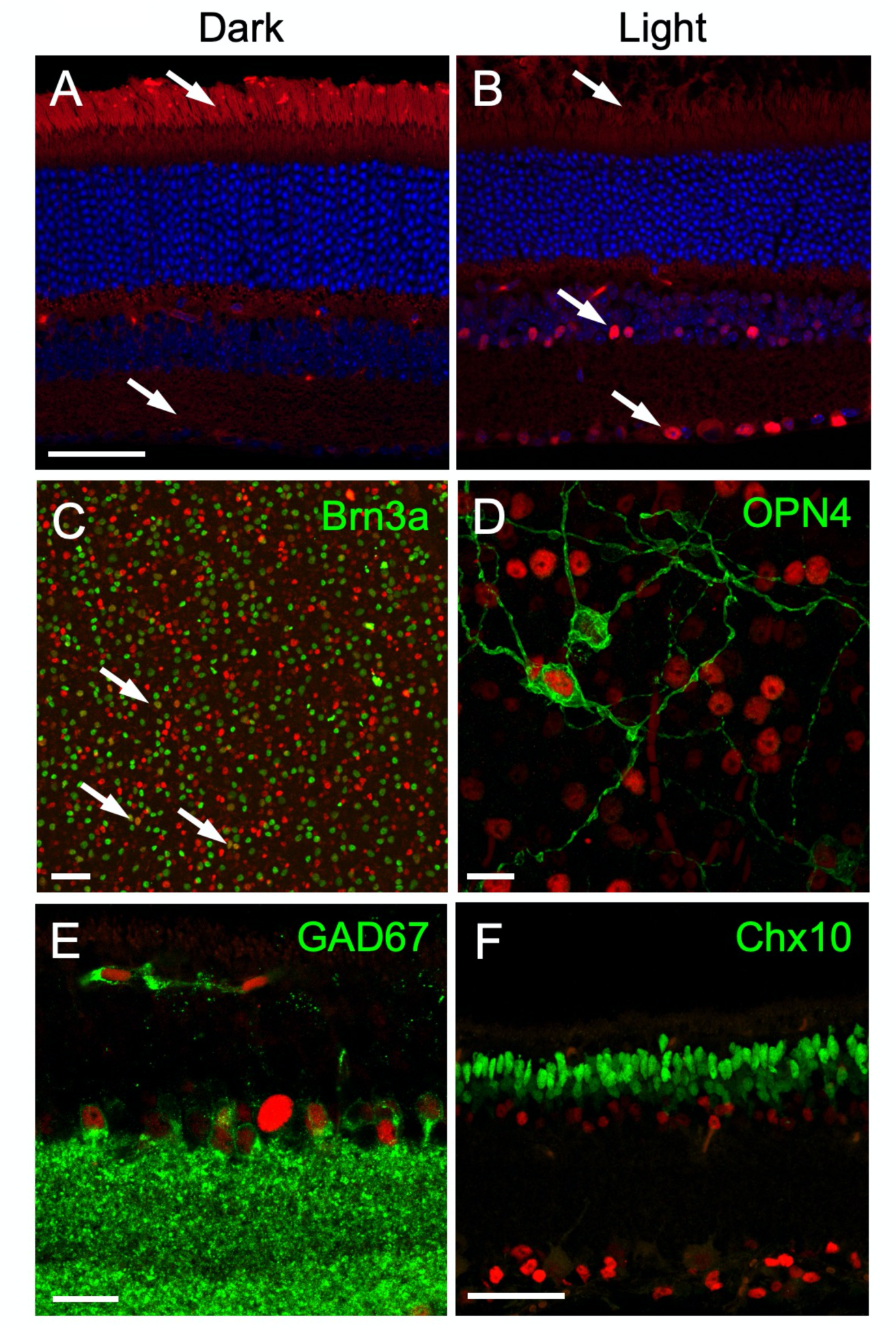
Retinal c-Fos responses to light occur amacrine and ganglion cells. Sections of wildtype retina stained for c-Fos in light **(A)** and dark **(B).** Arrows indicate outer segments which are c-Fos positive in the dark, and inner nuclear layer and ganglion cell layer cells that are c-Fos positive in the light. Scale bar 50µm. **(C)** whole mount retina labelled for Brn3a (ganglion cell marker) in green and c- Fos in red, arrow indicates co-labelled cells. Scale bar 50µm**. (D)** Whole mount retina labelled for opn4 (intrinsically photosensitive ganglion cells) in green and c-Fos in red. Scale bar 20µm. **(E)** Section of retina showing inner nuclear layer cells labelled for Gad67 (green) (GABAergic amacrine cells) some of which express c-Fos in response to light. Scale bar 20µm (**F**) Section of retina stained for bipolar cell marker Chx10, Light induced c-Fos (red) is not found colocalised with Chx10. Scale bar 50µm

**Fig. S2.**
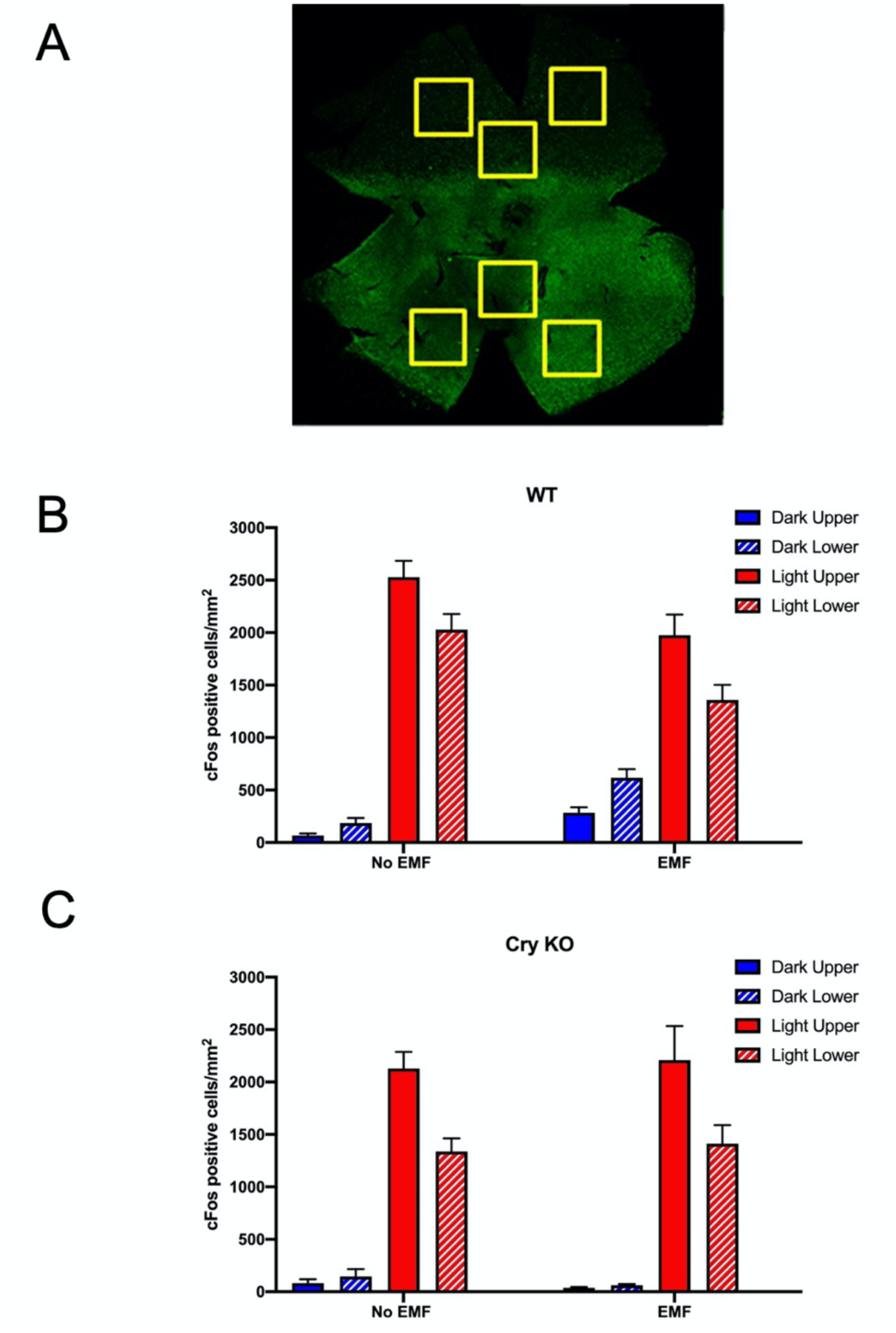
Retinal Location, Upper (dorsal) versus Lower (ventral). c-Fos effects in response to light and magnetic fields. Diagram of sampling of the retina for c-Fos quantification, 3 non-overlapping areas of the dorsal and ventral regions of each retina were imaged at low magnification (x10 objective, image size ∼0.5mm^2^). **(A)** c-Fos quantification in upper and lower quadrants of the wildtype retina, the lower retina shows significantly lower levels of light induced c-Fos, and there is a significant modulation by EMF. In the 3-way ANOVA (factors: EMF; Light; retinal location) significant factors are Light (F_1,38_=339.7, P<0.0001), Light x EMF interaction (F_1,38_=30.97, P<0.0001) and Retinal Location x Light interaction (F_1,38_=72.68, P<0.0001). **(B)** c-Fos quantification in upper and lower quadrants of the CRY-deficient mice, *Cry1^-/-^/Cry2^-/-^*. The lower retina again shows significantly lower levels of light induced c-Fos, but there is no modulation by EMF. In the 3-way ANOVA (factors: EMF; Light; retinal location) significant factors are Light (F_1,20_=279.8, P<0.0001), retinal location (F_1,20_=44.67, P<0.0001, and Retinal Location x Light interaction (F_1,20_=115.4, P<0.0001). Values plotted are means ±SEM.

**Fig. S3.**
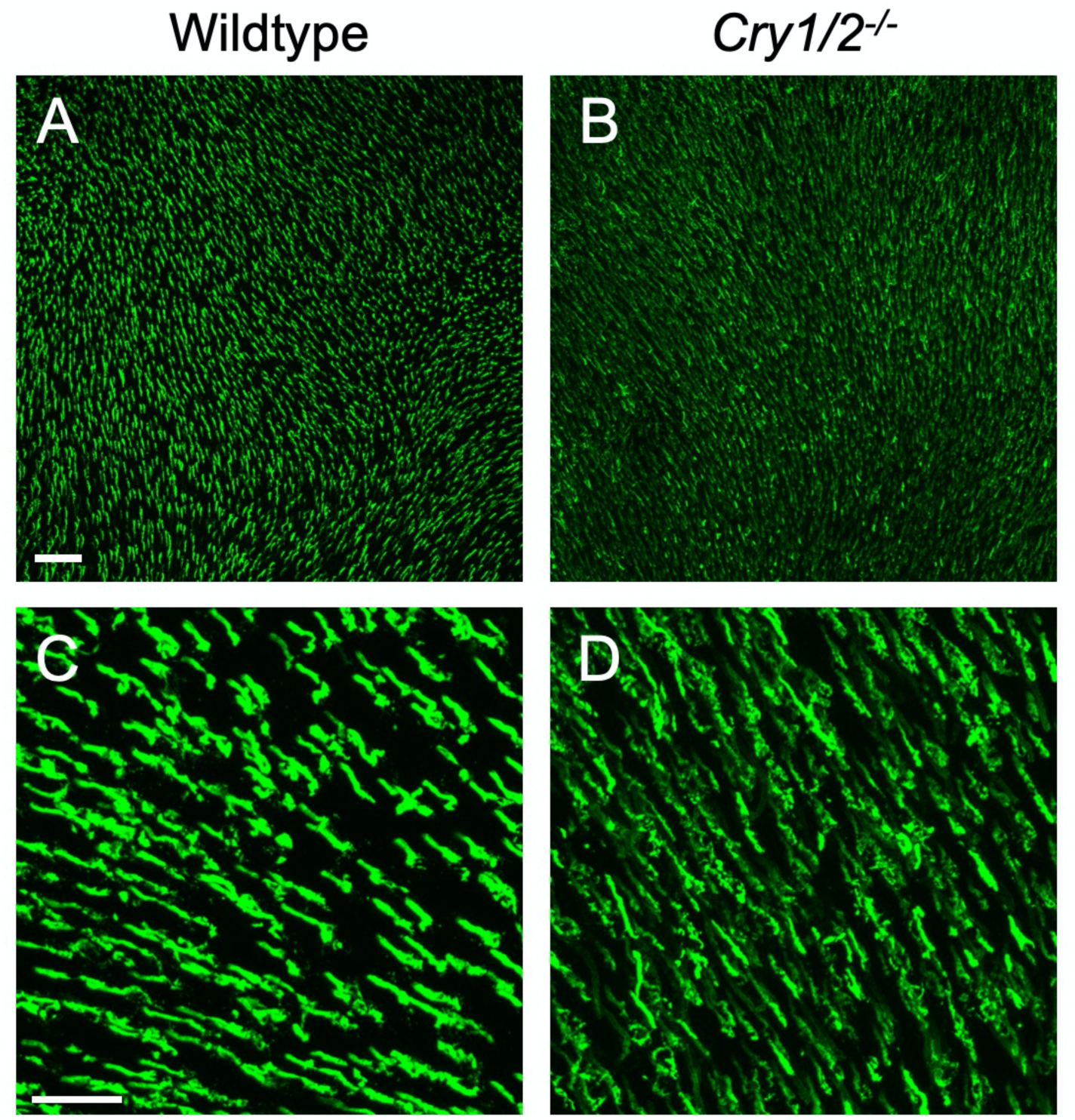
Changes in retinal photoreceptor morphology in Cry-deficient mice. Images of cones labelled with UVS opsin in the lower, ventral region of the retina. Retina from wildtype (**A**) and (**C**) and Cry deficient (**B**) and (**D**) mice. At higher magnification, (**C**) and (**D**) some minor deficits in the organisation of the cone outer segments were apparent in the Cry-deficient mice. Scale bar for A and B shown in A 50µm. Scale bar for C and D shown in C 20µm.

**Fig. S4.**
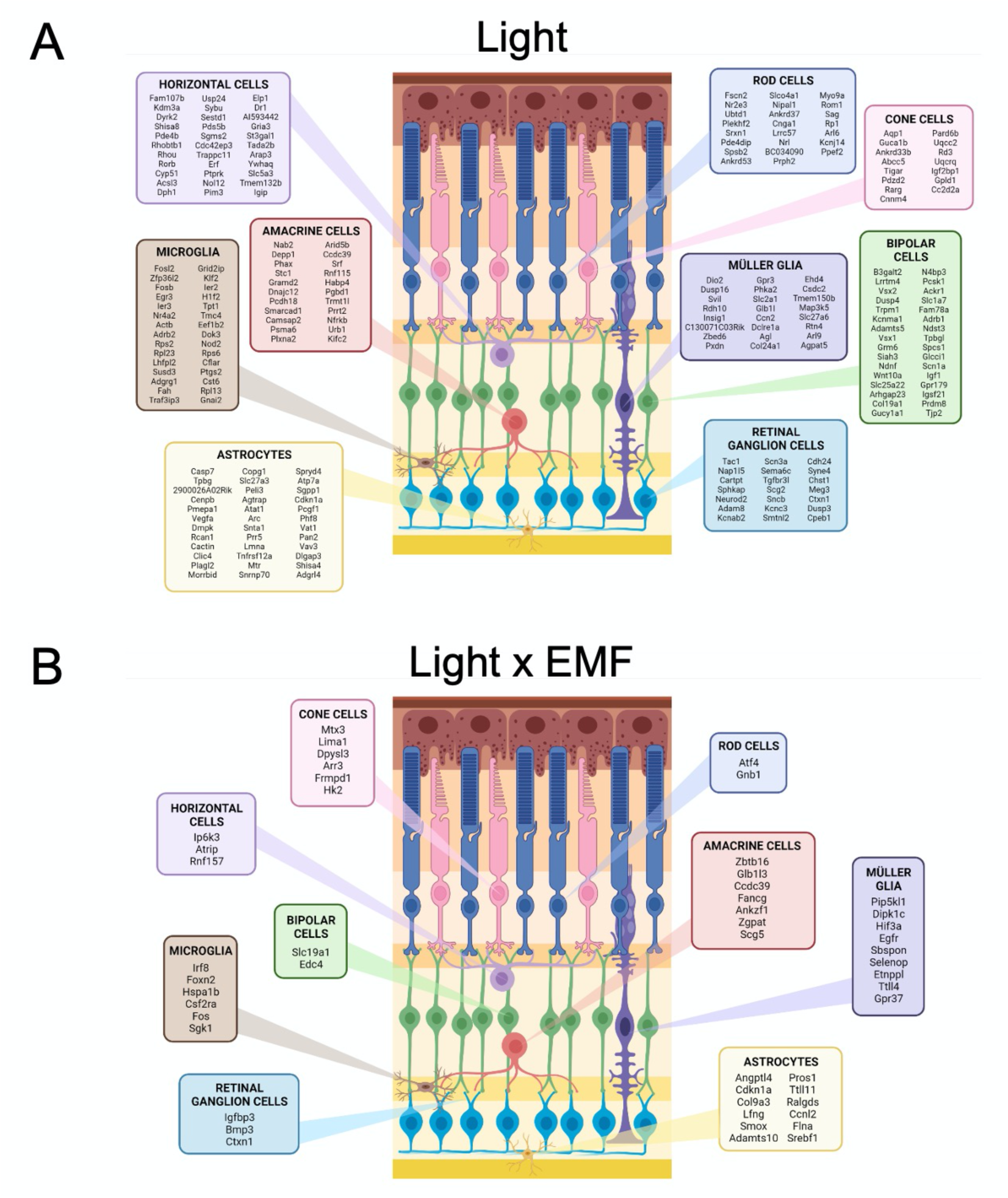
Mapping of cellular response of retinal gene expression in response to light and light x EMF. A schematic diagram of cell types in a section of retina, showing all the differentially expressed genes that map to a specific retinal cell type in response to (A) light and (B) light x EMF interaction.

**Fig. S5.**
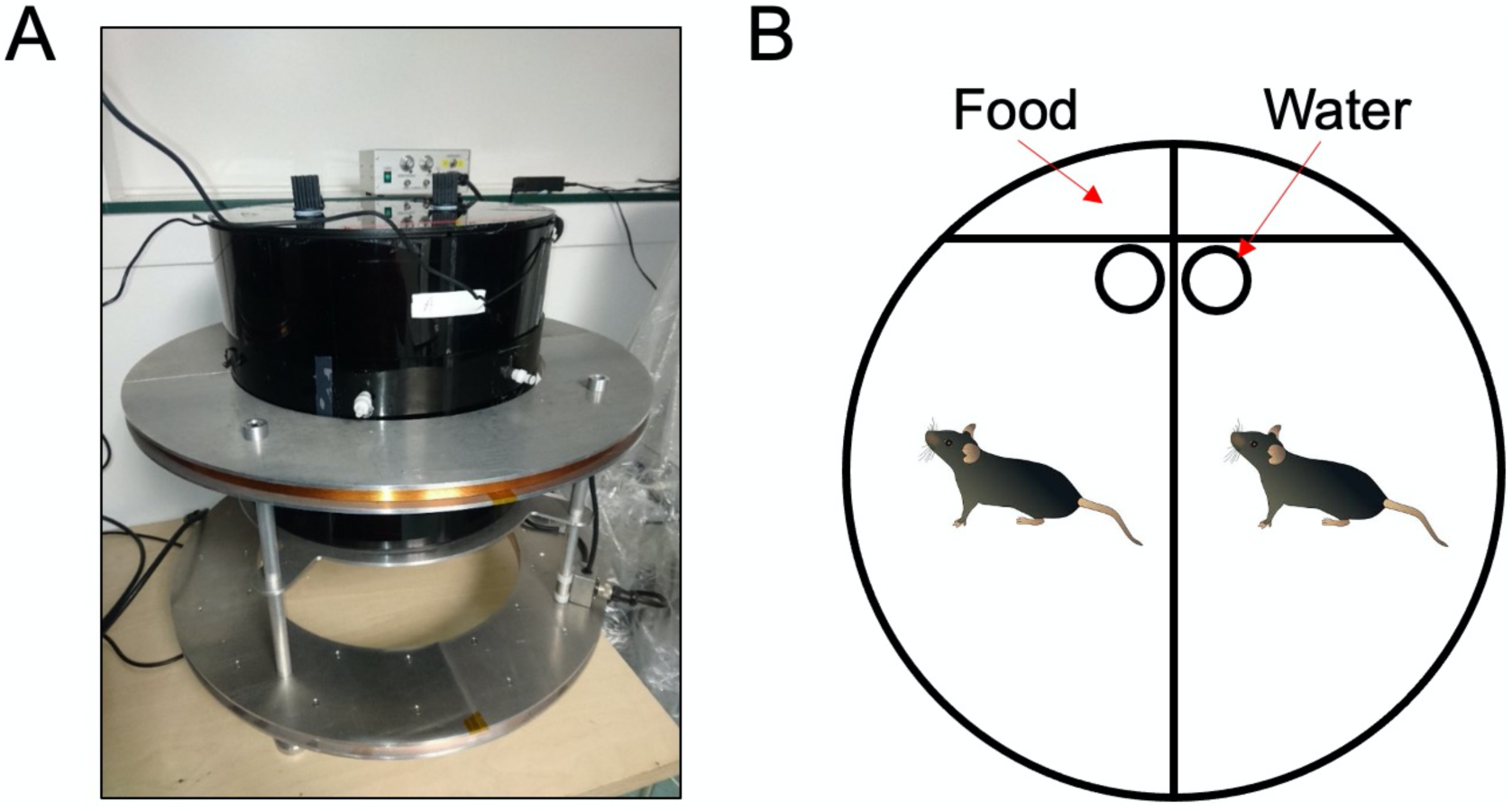
Magnetic field exposure system. **(A)** The custom made bifilar wound Helmholtz coil (diameter 50cm). When active, this coil generated a field strength of 100μT, or 0µT in the ‘no EMF’ condition in which electricity was still passed through the coil but no magnetic field was generated. Inside the Helmoltz coil is a black circular Perspex exposure chamber. **(B)** A schematic of the two separated housing zones within the EMF exposure chamber.

